# Effects of meglumine antimoniate and allopurinol treatment on the fecal microbiome profile in dogs with leishmaniosis

**DOI:** 10.1101/2024.12.27.630535

**Authors:** Joan Martí-Carreras, Marina Carrasco, Marc Noguera-Julian, Olga Francino, Rodolfo Oliveira Leal, Lluís Ferrer, Gaetano Oliva, Jenifer Molina, Xavier Roura

**Affiliations:** Nano1Health S.L. (N1H), Edifici EUREKA, Parc de Recerca UAB, Bellaterra, 08193 Barcelona, Spain; Molecular Genetics Veterinary Service (SVGM), Veterinary School, Universitat Autònoma de Barcelona, Barcelona, Spain; CIISA – Centre for Interdisciplinary Research in Animal Health, Faculty of Veterinary Medicine, University of Lisbon/Associate Laboratory for Animal and Veterinary Sciences (AL4AnimalS); Departament de Medicina i Cirurgia Animals, Universitat Autònoma de Barcelona, Bellaterra, 08193 Barcelona, Spain; Department of Veterinary Medicine and Animal Production, University of Naples Federico II, Naples, Italy; Nestlé Purina PetCare EUROPE, Purina Studios, Carrer Clara Campoamor, 2, Esplugues de Llobregat, Barcelona, Spain; Hospital Clínic Veterinari, Universitat Autònoma de Barcelona, Bellaterra, 08193 Barcelona, Spain

**Keywords:** Canine, gut microbiota, *Leishmania*, antimonial, dysbiosis

## Abstract

The combination of meglumine antimoniate and allopurinol is considered one of the most effective treatments for canine leishmaniosis caused by *Leishmania infantum*. This study investigated the effects of this treatment on the gut microbiome of 10 dogs from Spain, Portugal, and Italy via fecal shotgun metagenomic sequencing over six months. Dogs were sampled at baseline (BL) and at one (M1) and six (M6) months post-treatment. The gut microbiome of *Leishmania*-infected dogs (BL) is dominated by *Prevotella*, *Collinsella*, *Bacteroides*, and *Blautia*, with individual variability being the primary determinant of microbiome composition. No significant changes in alpha diversity (Shannon index, gene number) or beta diversity (Bray-Curtis dissimilarity, UniFrac distance) were detected between pre- and post-treatment time points, suggesting that treatment with meglumine antimoniate and allopurinol does not disrupt the gut microbiota. Minor trends in taxonomic shifts were noted, with slight increases in *Bifidobacterium pseudocantenulatum*, *Collinsella tanakaei*, and *Slackia piriformis* after treatment, but these changes were not statistically significant after correction for multiple testing. Linear discriminant analysis and multivariable modeling confirmed that the microbial community structure was resilient to treatment effects. Individual-specific microbiome differences in diversity accounted for 52% of the observed variability, underscoring the personalized nature of the gut microbiota in dogs. Importantly, no adverse microbiome disruptions were detected, even with prolonged allopurinol use. This study highlights the robustness of the canine gut microbiome during antileishmanial therapy and highlights the use of meglumine antimoniate and allopurinol without compromising gut microbial diversity or health. Further studies with larger cohorts are recommended to confirm these findings and explore the functional roles of the gut microbiota in modulating immune responses in *Leishmania*-infected dogs.

## Introduction

Canine leishmaniosis is caused by *Leishmania infantum*, a parasitic protozoan transmitted by the bite of phlebotomine sandflies (Paltrinieri et al., 2010). The combination of N-methyl-glucamine (meglumine antimoniate) and allopurinol is considered the first-line and most efficacious treatment for canine leishmaniosis (Oliva et al., 2010; Solano-Gallego et al., 2011). The most frequent side effects described with the use of meglumine antimoniate in dogs are pain and swelling at the injection site, fever, diarrhea, loss of appetite and azotemia (Denerolle & Bourdoiseau, 1999; Digiaro et al., 2024; Noli & Auxilia, 2005; Oliva et al., 2010; Roura et al., 2021; Slappendel & Teske, 1997). Allopurinol has also been associated with several associated effects, mainly urinary, including xanthinuria, kidney mineralization and urolithiasis (Jesus et al., 2022; Torres et al., 2016). Moreover, owing to the overuse of such compounds, genetic bases for antileishmanial drug resistance have been positively selected (Martí-Carreras et al., 2022), potentially decreasing drug efficacy.

Gut microbiome alteration can be related to diseases or drug side effects, which can change the functional composition and metabolic activities of bacteria in the gut and hence overall health status (Relman, 2015). Gut microbiome alterations linked to *Leishmania* infection have been previously studied in a wide array of hosts with varying evidence. In mice, microbiome changes linked to *Leishmania* infection vary by breed, with the CsS/Dem breeds showing resistance and the OcB/Dem breeds being more sensitive (Mrázek et al., 2024). However, in Syrian hamsters, no significant microbiome alterations are observed (Olías-Molero et al., 2022; Passos et al., 2020). In dogs, *Leishmania* infection increases Proteobacteria compared to healthy counterparts (Meazzi et al., 2022), while in humans, changes in *Ruminococcaceae* and *Gastranaerophilales* occur with visceral leishmaniosis (Lappan et al., 2019). Non-antibiotic drugs may also affect microbiomes (Maier et al., 2018), yet studies on antileishmanial treatments are limited. Miltefosine showed no effect in Syrian hamsters’ microbiome (Olías-Molero et al., 2023), however, the effects of other antileishmanial treatments on the fecal microbiome remain unstudied.

Molecular tools are now the standard for microbiome analysis in dogs using feces, as it is the most accessible sample type in clinical settings (Costa & Weese, 2019; Suchodolski, 2022). DNA shotgun metagenomic technique aims to sequence extracted DNA in a sample without prior amplification by PCR based on genomic DNA fragmentation and then randomly sequenced on a high-throughput sequencer. Unfortunately, this approach has rarely been used to assess the fecal microbiota in veterinary medicine (Quince et al., 2017; Sung et al., 2023).

This clinical study aimed to identify whether the most recommended antileishmanial treatment in dogs does have an impact on fecal microbiome composition via shotgun sequencing approach.

## Materials and methods

### Ethics

Dogs with confirmed leishmaniosis and not yet under treatment were eligible for the study. No untreated sick dogs were included in the study as a control group as it was against veterinarian ethical practice. Dog owners’ written consent was obtained for fecal sampling and access to veterinarian records, if required. A basic questionnaire was given for dog lifestyle data.

### Study design

The study was international, multicentric, prospective, controlled, randomized, and blinded. The inclusion criteria for each dog consisted of an appropriate diagnosis of leishmaniosis based on clinical signs and clinicopathological findings compatible with leishmaniosis and confirmation by positive quantitative serology, such as the indirect fluorescent antibody technique (IFAT) or enzyme-linked immunosorbent assay (ELISA), and/or identification of *Leishmania* parasites by cytology, histopathology, or polymerase chain reaction (PCR). Furthermore, dogs were not under treatment for leishmaniosis at the time of their first visit.

Several parameters were recovered for each enrolled dog: country of origin (Spain, Portugal, or Italy), date of sampling, sex, neutered status, breed, age, weight (kg), lifestyle (indoor, outdoor, or mixed), living conditions (city or countryside), cohabiting status, diet (commercial, homemade, or mixed) (**Table 1**).

**Table 1.** Dog data that were included in the study. This table includes all dog-related data acquired for the study: sex, breed, age, neutered status, country of origin, lifestyle, habitat, possible cohabitation, and type of diet.

| Dog | Sex | Breed | Age (years) | Neutered | Origin | Lifestyle | Habitat | Cohabitation | Diet |
| --- | --- | --- | --- | --- | --- | --- | --- | --- | --- |
| Dog 1 | Male | Boxer | 1 | No | Spain | Indoors | City | No | Commercial |
| Dog 2 | Male | Dogue of Bordeaux | 8 | Yes | Portugal | Mixed | Countryside | No | Commercial |
| Dog 3 | Male | Crossbreed | 9 | Yes | Italy | Mixed | City | No | Mixed |
| Dog 4 | Female | Pointer | 11 | Yes | Spain | Mixed | Mixed | 2 cats | Commercial |
| Dog 5 | Male | Crossbreed | 5 | No | Italy | Mixed | Village | No | Mixed |
| Dog 6 | Male | Crossbreed | 1 | Yes | Spain | Indoors | City | No | Commercial |
| Dog 7 | Female | Crossbreed | 11 | Yes | Spain | Outdoors | Countryside | 2 dogs | Commercial |
| Dog 8 | Male | Golden Retriever | 11 | No | Portugal | Mixed | City | No | Commercial |
| Dog 9 | Male | Crossbreed | 11 | Yes | Portugal | Mixed | Village | No | Commercial |
| Dog 10 | Female | American Bully | 6 | Yes | Spain | Indoors | City | No | Commercial |

Leishmaniosis treatment regime was homogeneous during the study and was based on the combination of subcutaneous meglumine antimonate (Glucantime^®^, Boehringer Ingelheim) at a dose of 50mg/kg/12h for 30 days, and oral allopurinol at 10 mg/kg/12h for at least 6 months (Oliva et al., 2010).

Among all three centers, 10 dogs were recruited that fulfilled the enrollment requirements. At each of the control points at diagnosis (BL) and after one and six months of treatment (M1 and M6), fecal samples were collected along with extra data on leishmaniosis state, LeishVet clinical stage (Solano-Gallego et al., 2009), health comorbidities and other drug used unrelated to leishmaniosis (**Table 2**).

**Table 2.** Medical data per dog at the time of sampling. Each dog was sampled three times, and their leishmaniosis was graded based on LeishVet Stage (Solano-Gallego et al., 2009), their comorbidities and non-leishmaniosis treatments were recorded.

| Dog | Sampling | LeishVet Stage | Comorbidities | Other treatments |
| --- | --- | --- | --- | --- |
| Dog 1 | BL | IV | Meningitis | Prednisone |
|  | M1 | II | Meningitis | Prednisone |
|  | M6 | IV | Meningitis | Prednisone |
| Dog 2 | BL | II |  | Fenbendazole |
|  | M1 | II |  |  |
|  | M6 | II |  | Telmisartan |
| Dog 3 | BL | II |  |  |
|  | M1 | II |  |  |
|  | M6 | II |  |  |
| Dog 4 | BL | III | Ehrlichiosis | Doxycycline |
|  | M1 | I |  |  |
|  | M6 | I |  |  |
| Dog 5 | BL | II |  |  |
|  | M1 | II |  |  |
|  | M6 | II |  |  |
| Dog 6 | BL | III |  |  |
|  | M1 | II |  | Enalapril |
|  | M6 | 0-I |  | Enalapril |
| Dog 7 | BL | II |  |  |
|  | M1 | I |  |  |
|  | M6 | 0 |  |  |
| Dog 8 | BL | II | Hypothyroidism | Telmisartan and levothyroxine |
|  | M1 | II | Hypothyroidism | Telmisartan and levothyroxine |
|  | M6 | II | Hypothyroidism | Telmisartan and levothyroxine |
| Dog 9 | BL | II | Skin lesions | Fluralaner and prednisolone |
|  | M1 | II |  | Fluralaner |
|  | M6 | II |  |  |
| Dog 10 | BL | III |  |  |
|  | M1 | II-III |  |  |
|  | M6 | 0 |  |  |

### Microbiome DNA extraction from fecal samples

Fecal samples (stool or rectal swabs) were collected in ZymoBIOMICS DNA/RNA Shield Collection Tubes with swab (R1106, Zymo Research Corporation; Los Angeles, CA, USA) to conserve the sample until shipment to laboratory facilities. Total DNA was extracted with the ZymoBIOMICS DNA Miniprep Kit (D4300; Zymo Research Corporation; Los Angeles, CA, USA) following the manufacturer’s recommendations.

### DNA sequencing

The extracted DNA was submitted to Macrogen (Macrogen, South Korea) for whole DNA shotgun metagenomic sequencing via an Illumina HiSeq 150×2 bp instrument, obtaining a median depth of 10 million reads 150 bp paired end. Sequencing data was obtained in fastq format. The study sequencing data is available under NCBI BioProject PRJNA1199800 under BioSample records SAMN45888860, SAMN45892024 - SAMN45892052 and demultiplexed reads under SRA41423504-41423475.

### Sequence data analysis

Sequencing reads (fastq) were quality filtered and trimmed for indices and low-quality bases using Trimmomatic v0.39 (LEADING:30, TRAILING:30, MINLEN:75, SLIDINGWINDOW:30:20) (Bolger et al., 2014). Taxonomic classification of sequencing reads to taxa are analyzed by mapping them to marker genes with Metaphlan3 v201901b using the database HUMANN3 (Beghini et al., 2021). Gene Richness/Number was determined from alignment of high-quality reads using Bowtie2 v2.4.1 (Langmead & Salzberg, 2012) against the Integrated Gene Catalog (Li et al., 2014).

Taxonomy data were further analyzed with R v4.1.1 (R Core Team, 2019) and the packages phyloseq v1.38.0 (McMurdie & Holmes, 2013), vegan v2.6.4 (Oksanen et al., 2024), lefser v1.4.0 (Segata et al., 2011), maaslin v1.8.0 (Mallick et al., 2021), and r_statix v0.7.0 (Kassambara, 2023). Count data can be accessed by species, genera, and phyla in **Supplementary Tables**.

## Results

During 2021, 10 client-owned dogs sick of leishmaniosis met the inclusion criteria and were enrolled in the study at one of the three participating university veterinary teaching hospitals, four from Universitat Autònoma de Barcelona (Spain), four from Faculdade de Medicina Veterinária de Lisboa (Portugal) and two from Università di Napoli (Italy).

Seven were male (three entire and four neutered), and three were spayed female. The median age of the dogs enrolled was eight years (range one to 11 years). There were six breeds represented, including crossbreed (n = 5) and 1 each of the following breeds: American Bully, Boxer, Golden Retriever, Pointer and Dogue of Bordeaux.

### Microbiome composition of *Leishmania*-infected dogs

Taxonomic affiliation revealed a grand total of 198 different species divided into 82 different genera and 10 phyla for the 10 individuals during the six months of the study (**Supplementary Tables**). An overview of the most abundant genera and species is displayed per dog (n = 10) and time point (BL, M1 and M6) in **Figure 1**.

**Figure 1.**
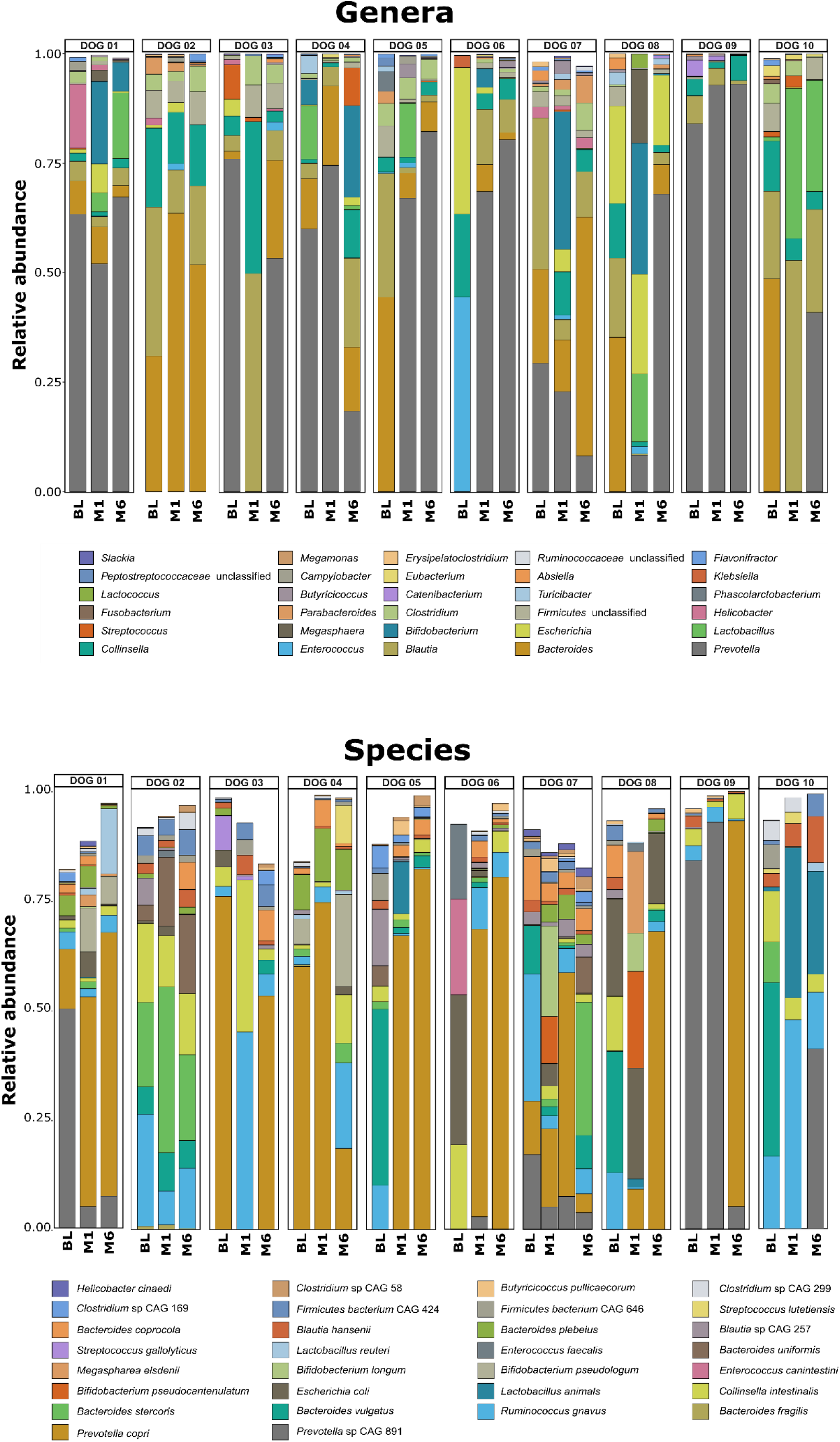
Relative abundance of bacteria at genus and species taxonomic levels. Stacked bar plots depicting the most abundant genera (upper panel) and species (bottom panel) by dog enrolled in the study (y-axis). Each taxon is colored independently, as shown in the legend. Each dog has three bar plots corresponding to pre-treatment (BL), and one- and six-months post treatment (M1 and M6).

Bacteria from the *Prevotellaceae* genus (mostly *Prevotella copri*) and the *Colinsella* genus (mostly *Colinsella intestinalis*) are dominant members of the canine microbiome in *Leishmania*-infected dogs together with *Bacterioides* (mainly *Bacteroides vulgatus* and *Bacteroides intestinalis*) and *Blautia* genera (**Figure 1**). The *Clostridium* genus is also present in many samples. Interestingly, *Colinsella intestinalis* shows lower abundance but higher prevalence than *Prevotella*. Similarly, *Ruminococcus gnavus* and *Bacteroides intestinalis* were frequently found in many of the samples (**Figure 1**). Notably, *Escherichia coli* was present in 33% of the samples, with no apparent association with any of the considered variables, as well as the *Enterococcus* genus, which is highly abundant in Dog06.

Among the most frequent bacteria genera (top 15), *Blautia* and *Bacteroides* genera are the dominant members of the canine microbiome in *Leishmania*-infected dogs, followed by *Prevotella* and *Colinsella* genera (**Figure 2a**). The *Clostridium* genus was present in many samples and dominant in sample Dog06. Several bacterial genera identified as members of *Firmicutes* phylum were also worthy of note. *Escherichia* genus was present in 33% of the samples, with no apparent association with any of the considered variables (**Figure 2a**). *Butyrucucicoccus*, *Flavonifractor*, *Absiella*, *Catenibacterium* and *Streptococcus* genera, together with several unclassified genera of *Peptostreptococcaceae* family correspond to minor genera in the gut microbiome. Nevertheless, their relative abundance per sample strongly varies between baseline samples, highlighting the individual differences in dog gut microbiome. Among these most prevalent genera, the most representative bacterial species (top 15) corresponded to *Ruminococcus gnavus* and *Collinsella intestinalis*, followed by *Blautia hensenii* and several uncultured *Firmicutes* bacteria (**Figure 2b**). *Escherichia coli*, *Bacteroides coprocola*, *Butyrucucicoccus pullicaecorum*, *Bacteroides sterconis*, *Colstriduim hiraconis, Flavonifractor plautii, Abisella dolichum* and *Bacteroides plebeius* are less represented species that are found in high abundance in some samples, such as *Escherichia coli*, which may correspond to dog specific microbiome composition.

**Figure 2.**
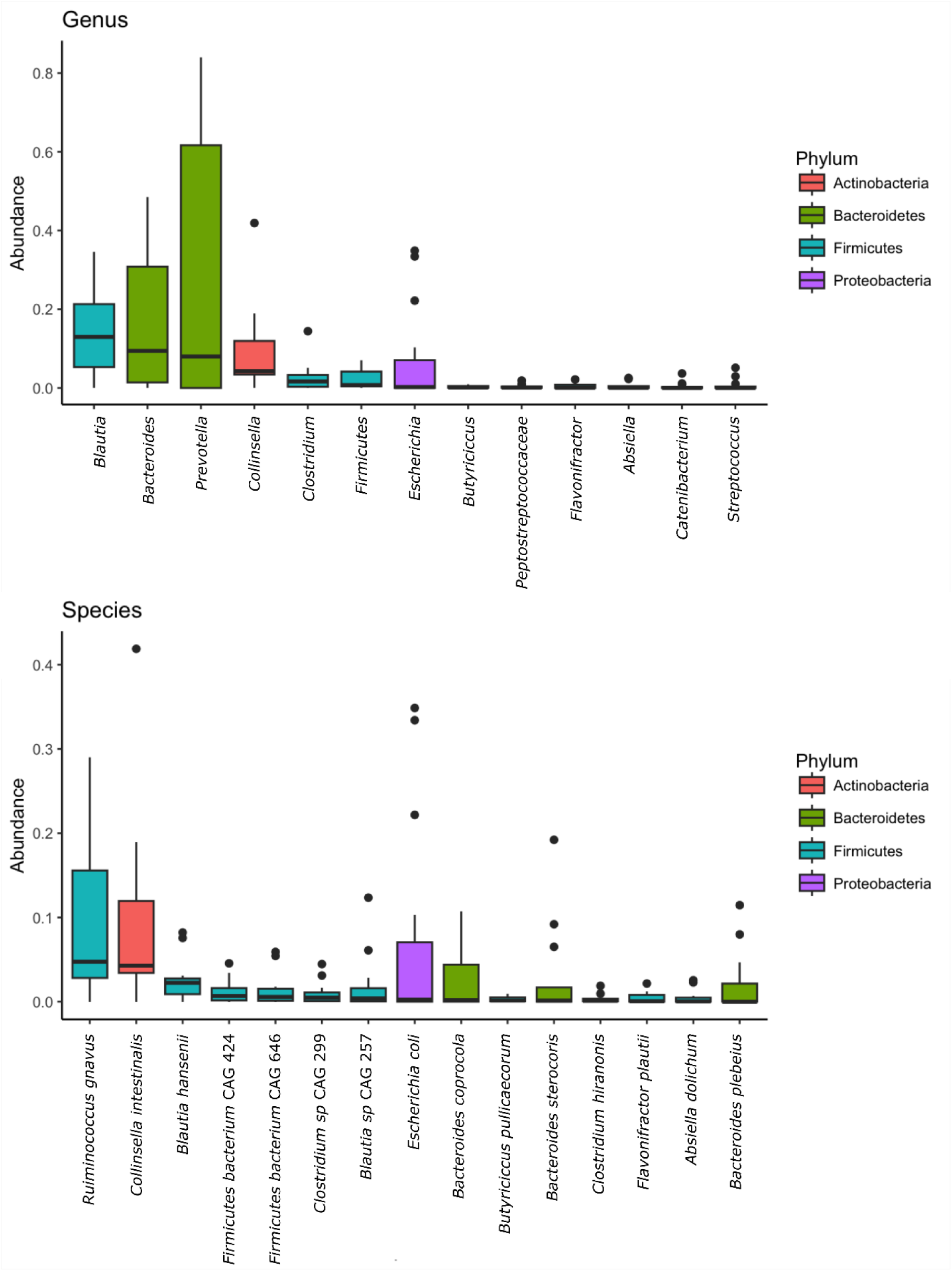
Summary of the relative abundance of the most prevalent bacteria in the study at genus and species taxonomic levels. The top panel shows the relative abundance (Y axis) of most prevalent bacteria at genus level (X axis). The bottom panel shows the relative abundance (Y axis) of most prevalent bacteria at species level (X axis). Phylum is indicated by color.

A comparison of the microbiome communities of each sample and dog by weighted UniFrac distance showed no clear grouping by non-metric multidimensional scaling (**Figure 3**). Bray-Curtis dissimilarities were statistically significant between individual dogs (PERMANOVA p-value = 0.001) but not between sampled timepoints (PERMANOVA p-value = 0.086), locations or any of the variables used in the study (**Table 1** and **Table 2**). The overall microbe structure depends more on the individual dogs (52% of the diversity) than on the timepoint. No overall differences in microbiome composition were detected for the duration of the study, as shown in **Figure 1**.

**Figure 3.**
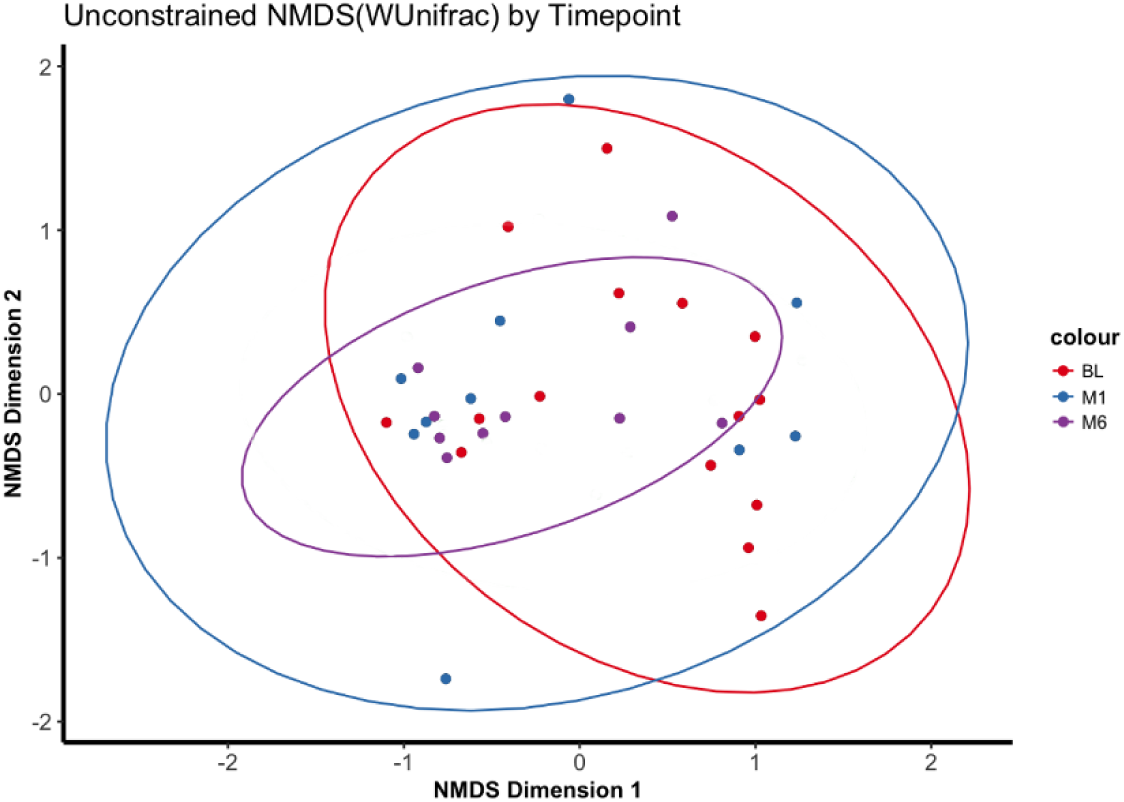
Projection of microbiome communities by non-metric dimensional scaling (NMDS) of each dog before and after treatment. The axis and units are arbitrary. Colors depict time point of sampling bar plots corresponding to pre-treatment (BL), and one- and six-months post-treatment (M1and M6).

### Microbiome alteration due to antileishmanial treatment

To determine whether the combined administration of meglumine antimoniate and allopurinol during the first month of treatment impacts the gut microbiome, samples were analyzed as pairs (BL vs M1). As shown in **Figure 4a**, there were no significant differences in taxonomy (Wilcox p-value = 0.36) or gene (Wilcox p-value = 0.92) diversity (alpha diversity) before treatment and after a month of combined therapy. A comparison between the microbiome communities before and after one month of treatment (BL vs M1) revealed no statistical difference due to the combined treatment for a month (PERMANOVA p-value = 0.426). Moreover, individual dog microbiome diversity represented 61% of the overall diversity.

**Figure 4.**
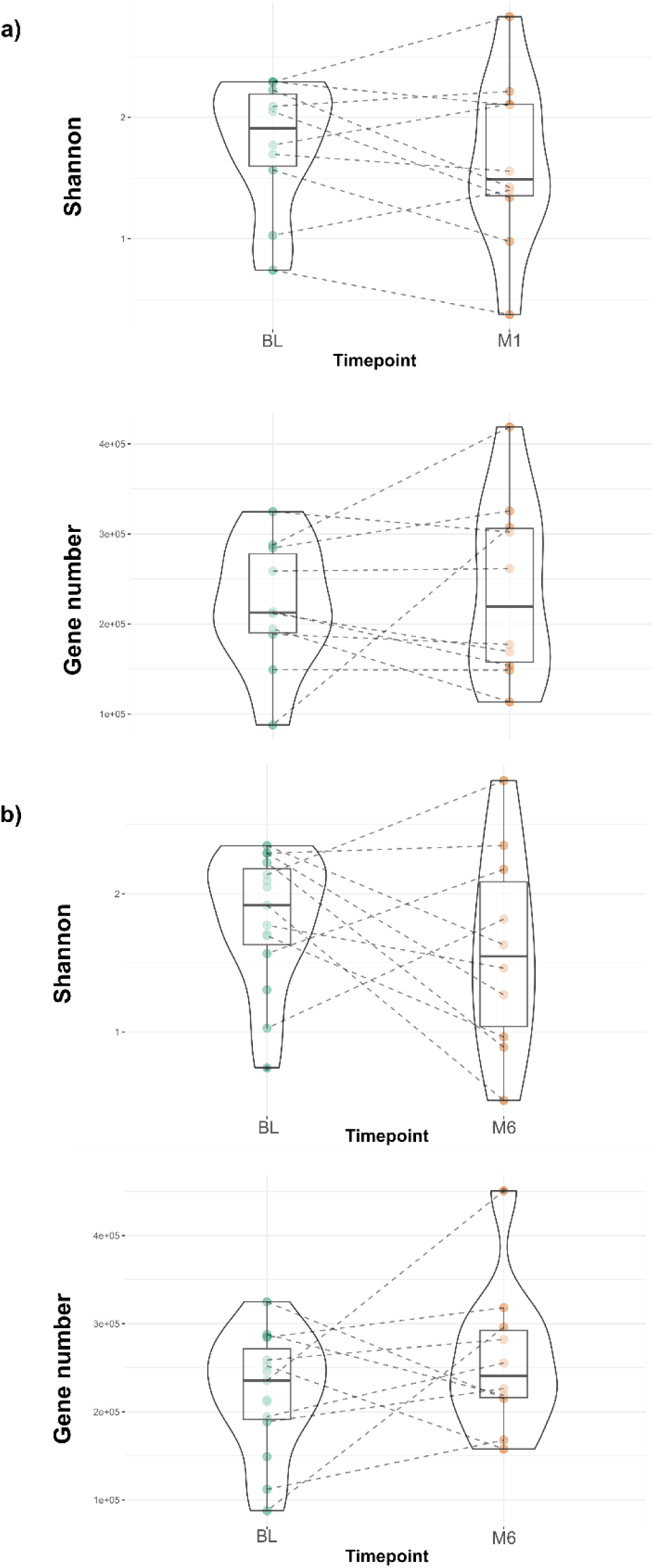
Comparison of taxonomic diversity and functional diversity before treatment and after one month of treatment. Panel a) Gut microbiome diversity is compared before (BL) and after one month of treatment (M1) for each individual using Shannon index (top panel) to estimate taxonomic diversity and gene number (bottom panel) to estimate functional diversity. Panel b) Gut microbiome diversity is compared before (BL) and after six months of treatment (M6) for each individual using Shannon index (top panel) to estimate taxonomic diversity and gene number (bottom panel) to estimate functional diversity.

Similar results were obtained after long-term administration of allopurinol, when samples before treatment and 6 months after initial treatment were analyzed as pairs (BL vs M6). As shown in **Figure 4b**, there were no significant differences in taxonomy (Wilcox p-value = 0.48) or gene (Wilcox p-value = 0.19) sample diversity (alpha diversity) before treatment and after six months of allopurinol treatment. A comparison of the microbiome communities (Bray-Curtis dissimilarity) before and after 6 months of treatment with allopurinol showed no statistical difference (PERMANOVA p-value = 0.11).

Individual species present in the M1 and M6 communities were compared with the same species present in the BL to identify differentially enriched or depleted taxa due to treatment. Linear discriminant analysis, using LEfSe (Segata et al., 2011), was performed, identifying possible enrichments of *Bifidobacterium pseudocantenulatum*, *Bifidobacterium longum*, *Collinsella tanakaei*, *Bifidobacterium pullorum* and *Allisonella histaminiformans* in M1 and *Prevotella copri* and *Slackia piriformis* in M6. The method detected a decrease in *Erysipelatoclostridium ramosum* in BL. After significance testing and correcting for multiple testing (Kruskal-Wallis with Bonferroni multiple testing correction, p-value > 0.005), no hits were statistically significant. Results were confirmed with microbiome multivariable association with linear models in MaAsLin (Mallick et al., 2021).

## Discussion

This study aimed to investigate the impact of antileishmanial treatment on the gut microbiome composition with 10 sick dogs of leishmaniosis from three clinical facilities (Spain, Italy, and Portugal). Treatment adherence and clinical follow up were compiled for the duration of the study (6 months). DNA extracted from fecal samples was sequenced using undirected metagenomics, an approach that allows the simultaneous detection of all DNA components of the microbiome (viruses, bacteria, archaea, or eukaryotes) compared with amplicon sequencing of marker genes (e.g., 16S) (Kim et al., 2024).

### Microbiome composition

The taxonomic composition of the gut microbiome in *Leishmania*-infected dogs over six months revealed remarkable individual variability, despite the presence of core taxa shared across the cohort. Across the 10 dogs sampled, the gut microbiome was dominated by members of the *Prevotella*, *Collinsella*, *Bacteroides*, and *Blautia* genera, which is consistent with previous studies indicating the role of these genera in canine gut health (Rojas et al., 2024; Suchodolski et al., 2008). The canine healthy gut bacteria composition is rich, strongly represented by *Prevotella*, *Fusobacterium*, *Clostridium* (sensu lato), *Lactobacillus*, *Streptococcus*, *Escherichia*/*Shigella*, *Blautia*, *Ruminococcus*, *Rombustia* and *Turicibacter* genera (Rojas et al., 2024; Suchodolski et al., 2008). Their relative abundance or presence in the fecal tract may depend on multiple factors, including age or feed (Rojas et al., 2024). As healthy dogs age, relative abundance of *Prevotella copri*, *Alloprevotella rava* and *Bacteroides coprocola* decrease, meanwhile *Escherichia coli* increases. Similarly, healthy dogs that do not consume kibble regularly have the relative abundances of *Bacteroides vulgatus*, *Caballeronia sordidicola*, *Enterococcus faecium*, *Erysipelatoclostridium ramosum*, *Blautia* sp. And *Clostridium* sp. increase, whereas that of healthy dogs that consume kibble regularly have an increase in the relative abundance of *Collinsella intestinalis*, *Turicibacter sanguinis*, *Megamonas funiformis*, *Holdemanella biformis*, and *Prevotella copri* (Rojas et al., 2024). In this study, such trends were not observed, as gut microbiome composition at pretreatment (BL) had no significant differential enrichment in bacterial relative abundance due to sex, breed, age, neutered status, country of origin, lifestyle, habitat, possible cohabitations, or type of diet (LEfSe and MaAsLin analysis). In fact, using Bray-Curtis dissimilarity (**Figure 3**), the primary driver of microbial diversity was the individual dog, accounting for 52% of the observed diversity. This finding is consistent with findings from both canine and human microbiome studies, where individual host factors (e.g., genetics, diet, and environment) have been shown to play a significant role in shaping microbial communities (Cuscó, Belanger, et al., 2017; Cuscó, Sánchez, et al., 2017; Knights et al., 2011).

In *Leishmania*-infected models, microbiome changes have been linked to *Leishmania*-infected mice (Mrázek et al., 2024). OcB/Dem breeds have been shown to be resistant to *Leishmania* infection, with positive associations with *Lactobacillaceae* and *Clostridiales*, *Peptococcaceae*, *Tannerellaceae* and *Burkholderiaceae*. Moreover, CsS/Dem breeds have been shown to be susceptible to *Leishmania* infections with positive associations with *Rikenellaceae*, *Deferribacteriaceae* and *Firmicutes* (Mrázek et al., 2024). These results do not translate to other rodent models, such as Syrian hamsters, where no significant microbiome alterations could be identified due to a *Leishmania* infection (Passos et al., 2020). Similarly, in *Leishmania*-infected dogs (sick or healthy) *Adlercreutzia*, *Prevotella*, *Clostridium*, *Enterococcus*, *Dorea*, *Megamonas*, *Roseburia* and *Escherichia* genera are differentially abundant in healthy dogs compared to *Leishmania*-infected dogs (Meazzi et al., 2022). In humans with visceral leishmaniosis, changes in microbiome composition have been reported in *Ruminococcaceae* and *Gastranaerophilales* between healthy and sick humans (Lappan et al., 2019). In this study, focusing on gut microbiome composition at pretreatment (BL), no members of *Peptococcaceae*, *Deferribacteriaceae*, *Adlercreutzia* or *Roseburia* were identified. The *Prevotella*, *Clostridium* and *Enterococcus* genera have many species present in the set of samples (40, 140 and 171 respectively), whereas the *Escherichia*, *Megamonas* or *Dorea* genera, despite being present and having several species representatives (20, 10 and 10 respectively) their species diversity is lower than *Prevotella*, *Clostridium* and *Enterococcus* genera. The relative abundance of these genera seems to be correlated with their diversity, having *Prevotella*, *Clostridium* and *Enterococcus* genera a higher relative abundance than *Escherichia*, *Megamonas* or *Dorea* genera.

Furthermore, pretreatment (BL), *Prevotella copri* and *Collinsella intestinalis* were prominent members of the gut microbiome, with the former being one of the most abundant species (**Figure 1 and Figure 2**). These genera are often linked to the digestion of complex carbohydrates and fibers, indicating their potential importance in maintaining gut homeostasis (Flint et al., 2012), even in the context of a chronic infectious disease such as leishmaniosis. Moreover, the dominance of *Blautia* and *Bacteroides* is notable because these genera are typically associated with anti-inflammatory properties and short-chain fatty acid (SCFA) production (Holmberg et al., 2024). The high prevalence of these taxa, particularly *Blautia*, which is linked to SCFA production, such as butyrate, suggests that even when dogs are infected with *Leishmania*, their gut microbiomes retain key microbial groups important for gut barrier function and immune modulation (Martin-Gallausiaux et al., 2021). Interestingly, several minor bacterial genera, including *Butyricicoccus*, *Flavonifractor*, *Absiella*, and *Catenibacterium*, presented high variability in relative abundance across samples (**Figure 1 and Figure 2**). These genera, particularly *Butyricicoccus* and *Flavonifractor*, are known SCFA producers, which play crucial roles in gut health by modulating the host immune response and maintaining intestinal barrier function (Louis & Flint, 2017). While their presence was not significant enough to drive large-scale microbiome changes, their sporadic occurrence in certain dogs might reflect individual differences in gut physiology or the immune response to *Leishmania* infection. Notably, that individual changes in the gut microbiome composition of specific taxa might not have a relevant impact on health status as a diverse microbiome does entail good functional redundancy, where the gut microbiome maintains its overall metabolic potential despite changes in microbial composition (Moya & Ferrer, 2016). *Collinsella intestinalis*, while not as abundant as *Prevotella*, displayed a higher prevalence, suggesting that this taxon may have a more consistent, albeit lower-level, role in the microbial community across different dogs, possibly contributing to maintaining metabolic functions over time, even in the presence of infection. The presence of *Escherichia coli* in 33% of the samples, without any clear association with the variables studied, is an intriguing finding. *Escherichia coli* is commonly associated with inflammation, especially in conditions such as inflammatory bowel disease in humans (Dubinsky et al., 2022). However, in this case, its sporadic appearance suggests that it may not play a central role in the microbiome changes associated with *Leishmania* infection or antileishmanial treatment, possibly reflecting individual variations in gut health or transient colonization by opportunistic pathogens.

Overall, these findings suggest that while *Leishmania* infection and its treatment do not dramatically alter the gut microbiome of infected dogs, there is substantial individual variability in microbiome composition. This variability may explain the observed resilience of the gut microbiome to infection and treatment and aligns with the increasing understanding of individual-specific microbiome responses in chronic diseases (Shoaie et al., 2015). Future studies should explore the functional consequences of these microbial dynamics, particularly the role of SCFA-producing bacteria in modulating the immune response during chronic *Leishmania* infection.

### Microbiome alteration due to antileishmanial treatment

The comparative analysis between baseline (BL) and post-treatment time points (M1 and M6) aimed to identify any significant shifts in microbiome composition due to the administration of meglumine antimoniate and allopurinol (M1) and long-term administration of allopurinol (M6). Our results indicate that the overall microbial diversity, both alpha diversity (Shannon index and gene number) and beta diversity (Bray-Curtis dissimilarity and weighted UniFrac), remained largely unchanged throughout the study.

Although the number of dogs included in the present study was not very large, the absence of statistically significant microbiome changes over time (p > 0.05) suggests that the gut microbiome changes in these *Leishmania*-infected dogs could not be linked to any common cause during the study. These findings suggest that the combined therapy (M1) or long-term single therapy (M6) does not induce significant changes in the microbiome communities during treatment.

Small enrichment trends, albeit not significant, were identified in M1 for *Bifidobacterium pseudocantenulatum*, *Bifidobacterium longum*, *Collinsella tanakaei*, *Bifidobacterium pullorum* and *Allisonella histaminiformans* and *Prevotella copri* and *Slackia piriformis* in M6. Small depletion trends, albeit not significant, were identified in the BL treatment for *Erysipelatoclostridium ramosum.* These results could be interpreted either as false positives from LefSe, as hits disappeared after multiple testing correction and were not present in MaAsLin2 analysis, or hits with low statistical power due to the small sample size.

The microbiome communities seem to be unaffected by the given treatment, with only uncertain evidence indicating small enrichments in *Bifidobacterium*, *Collinsella*, *Allisonella*, *Prevotella*, and *Slackia* genera. These taxa are commonly found in healthy microbiomes, suggesting a possible neutral effect of the treatment to the gut microbiome composition, which may be crucial for maintaining gut health and overall homeostasis during disease and recovery (Kinross et al., 2011). Drugs used to treat *Leishmania*, such as antileishmanial agents, may have similar effects.

Non-antibiotic drugs have been shown to alter microbial communities, potentially disrupting the balance of the gut microbiome, although the exact mechanisms remain largely unknown (Maier et al., 2018). However, studies examining the direct impact of antileishmanial treatments on the microbiome are limited, highlighting the need for such studies. One recent study investigating the effects of miltefosine in *Leishmania* infantum-infected Syrian hamsters reported that the drug did not significantly alter the gut microbiome composition (Olías-Molero et al., 2023). These findings suggest that miltefosine may not impact microbial populations in this model, yet possible effects may exist for other drugs, such as meglumine antimoniate or allopurinol or in a different disease models or hosts, such as dogs or humans. In the present study, meglumine antimoniate and allopurinol treatment had no significant effect on microbiome populations before and after treatment in a relevant model and host (dogs) with clear implications for leishmaniosis treatment and *Leishmania* management. This study promotes that standard antileishmanial treatments can be administered without major concerns regarding the disruption of gut microbiome, which is critical for maintaining the overall health of infected dogs. However, owing to the significant individual variability observed in the microbiome composition, individual treatment plans, such as dietary changes or the use of probiotics may help during the treatment and recovery periods of dogs with leishmaniosis, as previously recommended (Solano-Gallego et al., 2011).

## Conclusion

In conclusion, this study provides evidence that the recommended combined treatment of meglumine antimoniate and allopurinol during the first six months does not significantly alter the gut microbiome composition of dogs with leishmaniosis. Moreover, most of the observed changes in the gut microbiome are individual specific, highlighting the unique and dynamic nature of microbiome gut communities in dogs. Owing to the low degree of possible enrollment due to the study constraints, further research should be considered to discard any minor change or contribution during the beforementioned treatment.

## Acknowledgments

This study was sponsored by Nestlé Purina PetCare EUROPE.

